# Unveiling Vertebrate Development Dynamics in Frog *Xenopus laevis* using Micro-CT Imaging

**DOI:** 10.1101/2024.06.11.598452

**Authors:** Laznovsky Jakub, Kavkova Michaela, Reis Alice, Robovska-Havelkova Pavla, Krivanek Jan, Zikmund Tomas, Kaiser Jozef, Buchtova Marcela, Harnos Jakub

## Abstract

**Background:** *Xenopus laevis*, the African clawed frog, is a versatile vertebrate model organism employed across various biological disciplines, prominently in developmental biology to elucidate the intricate processes underpinning body plan reorganization during metamorphosis. Despite its widespread utility, a notable gap exists in the availability of comprehensive datasets encompassing *Xenopus’* late developmental stages.

**Findings:** In the present study, we harnessed micro-computed tomography (micro-CT), a non-invasive 3D imaging technique utilizing X-rays to examine structures at a micrometer scale, to investigate the developmental dynamics and morphological changes of this crucial vertebrate model. Our approach involved generating high-resolution images and computed 3D models of developing *Xenopus* specimens, spanning from premetamorphosis tadpoles to fully mature adult frogs. This extensive dataset enhances our understanding of vertebrate development and is adaptable for various analyses. For instance, we conducted a thorough examination, analyzing body size, shape, and morphological features, with a specific emphasis on skeletogenesis, teeth, and organs like the brain at different stages. Our analysis yielded valuable insights into the morphological changes and structure dynamics in 3D space during *Xenopus’* development, some of which were not previously documented in such meticulous detail. This implies that our datasets effectively capture and thoroughly examine *Xenopus* specimens. Thus, these datasets hold the solid potential for additional morphological and morphometric analyses, including individual segmentation of both hard and soft tissue elements within *Xenopus*.

**Conclusions:** Our repository of micro-CT scans represents a significant resource that can enhance our understanding of *Xenopus’* development and the associated morphological changes. The widespread utility of this amphibian species, coupled with the exceptional quality of our scans, which encompass a comprehensive series of developmental stages, opens up extensive opportunities for their broader research application. Moreover, these scans have the potential for use in virtual reality, 3D printing, and educational contexts, further expanding their value and impact.

**Graphical abstract & lay summary:** 3D images of selected developmental stages of *X. laevis* in a comparison (scale bar = 10 mm).

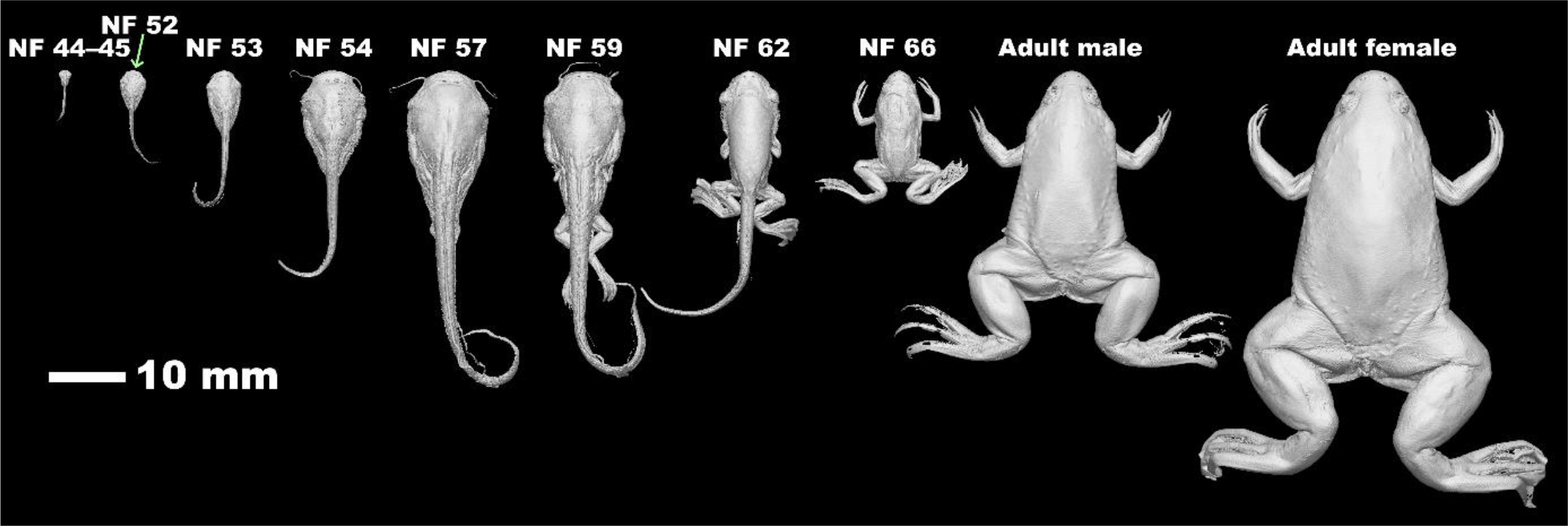

**Lay summary:** X-ray tomography was used to examine the African clawed frog (*Xenopus laevis*). An extensive data set of specimens from tadpoles to adult frogs provides novel insights into the changes and developmental dynamics of selected structures, which opens avenues to an improved understanding of this crucial animal model.

## Background

*Xenopus laevis*, commonly known as the African clawed frog, serves as a fundamental model organism in the field of life sciences. Its widespread adoption particularly in biological research can be attributed to the ease of breeding and housing, as well as the substantial size and manipulability of its eggs and embryos^1,2^. Over the years, this species has been extensively investigated across various disciplines, including genetics, embryology, and developmental, cell, and regenerative biology^3-5^. While initial attempts at anatomical descriptions of *X. laevis* date back a century^6^, subsequent studies have, to varying degrees, concentrated only on specific anatomical regions^7-25^. Despite these valuable contributions, many past approaches to describing frog anatomy failed to preserve the intricate morphological details, making them unsuitable for detailed and comparative studies. In response to these limitations, a detailed description of an adult male frog’s anatomy using a non-destructive micro-CT method was recently provided^7^. Nevertheless, the spatiotemporal dynamics of *X. laevis’* late development, along with the comparison between adult male and female frogs, remain largely unexplored in sufficient detail.

To fill this knowledge gap and gain a more comprehensive understanding of *X. laevis’* late development, we employed micro-CT, a powerful imaging technology for examining small objects at the micrometer scale^26^. Micro-CT utilizes X-rays to generate high-resolution 3D images, making it an ideal tool for investigating developmental and morphological changes, particularly in small embryos or tadpoles^26^. In principle, micro-CT imaging is based on capturing a series of 2D X-ray radiographs from various angles and then mathematically processing them through tomographic reconstruction, resulting in a 3D matrix representing volume density. An advantage of micro-CT lies in its ability to image bones and, when combined with various contrast methods, also soft tissues and blood vessels within the same sample^26^.

Through micro-CT, we present high-resolution images and computed 3D models of developing *X. laevis* tadpoles, froglets, and adult frogs, which offer great potential for research fields like developmental and comparative biology. Additionally, we conducted preliminary analyses of this dataset to illustrate its versatility. Specifically, our preliminary examination covered morphological changes in body size, shape, and skeletogenesis from selected premetamorphosis, prometamorphosis, and climax metamorphosis stages through the froglet stage up to adulthood. Our findings unveil novel insights into the intricate morphological dynamics of late *X. laevis’s* development, providing unprecedented levels of detail. This research contributes a unique dataset concerning the developmental and morphological changes in *X. laevis*, including the adult stages, shedding light on the dynamics of vertebrate development, and bearing broader implications for future analyses of *Xenopus*.

### Sampling Strategy

In our micro-CT study of *Xenopus*, we employed a careful sampling strategy to encompass the entire developmental spectrum of this amphibian species. The selection of developmental stages was based on specific morphological features and markers that serve as critical indicators of *X. laevis’* development. By strategically choosing nine distinct developmental stages, we aimed to capture the comprehensive process of morphogenesis, spanning from premetamorphosis tadpoles to fully mature adult frogs. These stages were carefully chosen with reference to specific criteria, such as the presence and dimensions of legs and the tail, or the rearrangements of the head, as outlined in the tables of Nieuwkoop and Faber^27^, and Zahn and colleagues^28^.

Our sampling included the following crucial stages of *X. laevis’* development: **Premetamorphosis** (stages 44-45, 52, and 53, according to Nieuwkoop and Faber, NF): These early stages provide insight into the initial stages of development, as *Xenopus* transforms from tadpole to froglets.

**Prometamorphosis** (NF stages 54 and 57): This phase represents an intermediate stage where significant changes in the head, limb buds, and tail are occurring, leading slowly towards the final metamorphosis.

**Climax Metamorphosis** (NF stages 59 and 62): These stages are characterized by the most profound changes, marking the peak of metamorphosis.

**Froglets** (NF stage 66): Froglets represent a transitional stage, bridging the gap between tadpoles and fully mature adult frogs.

**Adult Male** and **Female Frogs**: These adult stages represent the endpoint of *Xenopus* development, allowing us to observe fully mature individuals and to compare differences between sexes.

Our selection of these key developmental stages was performed with regard to their significance in the overall developmental process of *X. laevis*. By examining these specific stages, we aimed to gain a comprehensive understanding of the morphological transformations and developmental milestones that occur during *X. laevis’* development, from its primary aquatic tadpole phase to its secondary aquatic adult frog stage^29^. This sampling strategy provided a detailed and holistic view of *X. laevis’* late development, enabling us to perform valuable analyses leading to insights and preliminary conclusions about this species’ morphological changes throughout its life cycle.

### Source of Samples

Our work with *X. laevis* adhered to Czech animal use and care laws and received approval from local authorities and committees (MSMT-30784/2022-1; Animal Care and Housing Approval: 45055/2020-MZE-18134, Ministry of Agriculture of the Czech Republic; and Animal Experiments Approval: CZ 62760214, State Veterinary Administration/Section for South Moravian Region). *Xenopus* embryos were generated and cultivated following standard protocols. Briefly, testes were surgically removed from anesthetized males (20% MS-222, Sigma-Aldrich, A5040) and transferred to cold 1x Marc’s Modified Ringers (MMR; 100mM NaCl, 2mM KCl, 1mM MgSO_4_, 2mM CaCl_2_, 5mM HEPES, buffered to pH 7.4), supplemented with 50 μg/mL of gentamycin (Sigma-Aldrich, G3632). To induce egg laying, fully mature *Xenopus* females were injected with 260 U of human chorionic gonadotropin (Merck, Ovitrelle 250G) into the dorsal lymph sac approximately 12-16 hours before use and were kept overnight at 18°C. For fertilization, eggs were extracted from induced females directly into a Petri dish and mixed with a piece of testes in 0.1x MMR at 18-21°C. The subsequent development of *Xenopus* tadpoles, froglets, and adult frogs adhered to the approved cultivation protocols (see above). At a designated time point, *Xenopus* specimens were anesthetized and fixed in a buffered 4% paraformaldehyde solution (1004965000, Merck) for 3 hours (tadpoles) or overnight (froglets and adult frogs), and staged according to the tables of Nieuwkoop and Faber^27^, and Zahn and colleagues^28^. Real images of *Xenopus* specimens used for micro-CT analysis are presented in **Suppl. Fig. 1**.

### Micro-CT Scanning

Prior to scanning, samples were placed in either a 2 ml Eppendorf tube, a 15 ml/50 ml Falcon tube (depending on sample size), or in the case of the adult frogs, a 500 ml plastic container. To prevent the motion and drying of the sample during the micro-CT scan, all samples were mounted in a 1% agarose gel (Top-BIO, P045). Micro-CT measurements were conducted using the GE Phoenix v|tome|x L 240 laboratory system, which is equipped with a 180kV/15W nanofocus X-ray tube and a 4000 x 4000-pixel flat panel detector with a 100 μm pixel size. Scan conditions for each stage are summarized in **Suppl. Table 1**. The tomographic reconstruction was performed using the GE Phoenix datos|x 2.0 software. The reconstructed data were imported into VG Studio MAX 2022.1 for data analysis and visualization.

All *Xenopus* samples were initially scanned in their native state to visualize bone structures. Subsequently, specimens were stained with 1% iodine (Penta, 21210-11000) in a 90% methanol (Penta, 17570-30250) solution to enhance the contrast and visualize soft tissues. As for teeth shown in Fig. 3, the separated jaws were scanned and stained. The dehydration and staining times for each *Xenopus* developmental stage are detailed in **Suppl. Table 2** (for adult frogs, the staining solution was changed three times, as indicated in the table).

### Data Quality Control and Limitations

All analyzed samples, comprising tadpoles, froglets, and adult frogs, were uniformly preserved and handled before scanning. The primary variable affecting data quality was the varied voxel size of each dataset, arising from different sample sizes. The smallest sample, a tadpole stage (NF 44-45), featured a voxel size of 5 μm, while the largest sample, an adult female frog, was scanned with a voxel size of 40 μm.

The voxel size difference is directly correlated with sample dimensions. The GE L240 X-ray source’s cone beam geometry and the detector’s field of view allowed smaller samples to be placed closer to the X-ray source, resulting in a smaller voxel size (higher resolution) due to cone beam magnification. Although samples exhibited variable voxel sizes, this variability does not necessarily limit data analysis. Larger samples, despite having larger voxel sizes, still enabled the recognition of structures analyzed, as these structures were proportionately larger in larger animals scanned with larger voxel sizes.

All *Xenopus* specimens underwent two sequential scans. Initially, native scans were performed, following the principle of micro-CT imaging of samples in their native form without staining, enabling the visualization of dense mineralized structures like bones and teeth. Subsequently, all scans were stained as described in **Suppl. Table 2**. To prevent movement and drying of samples during data acquisition, all samples were fixed in polypropylene conical tubes with 1% agarose gel. Throughout data acquisition, no observable sample shrinkage occurred. If any negligible shrinkage or drying did occur, we anticipate the volume reduction to be proportionally consistent across all *Xenopus* samples.

### Atlas of *X. laevis’* Late Development

In order to provide a more comprehensive understanding of *X. laevis’* development during metamorphosis and organogenesis, we first collected specimens representing the key frog stages, including premetamorphosis, prometamorphosis, climax metamorphosis, as well as froglets, and adult male and female frogs (see Sampling Strategy). Initially, we performed native sample scanning for hard tissue visualization and subsequently stained and scanned the samples for soft tissue visualization (see Micro-CT Scanning). As a result, we were able to collect and analyze all major stages of *X. laevis’* late development. The entire collection of the *Xenopus* atlas is presented in **Fig. 1**, and all raw data can be found in Zenodo repository (see section Data availability). Subsequently, with the atlas, we provide here several examples of specific analyses of *X. laevis’* late development, including an analysis of head development, teeth, bone growth dynamics, ossification, and brain development with its associated nerves.

**Figure 1.**
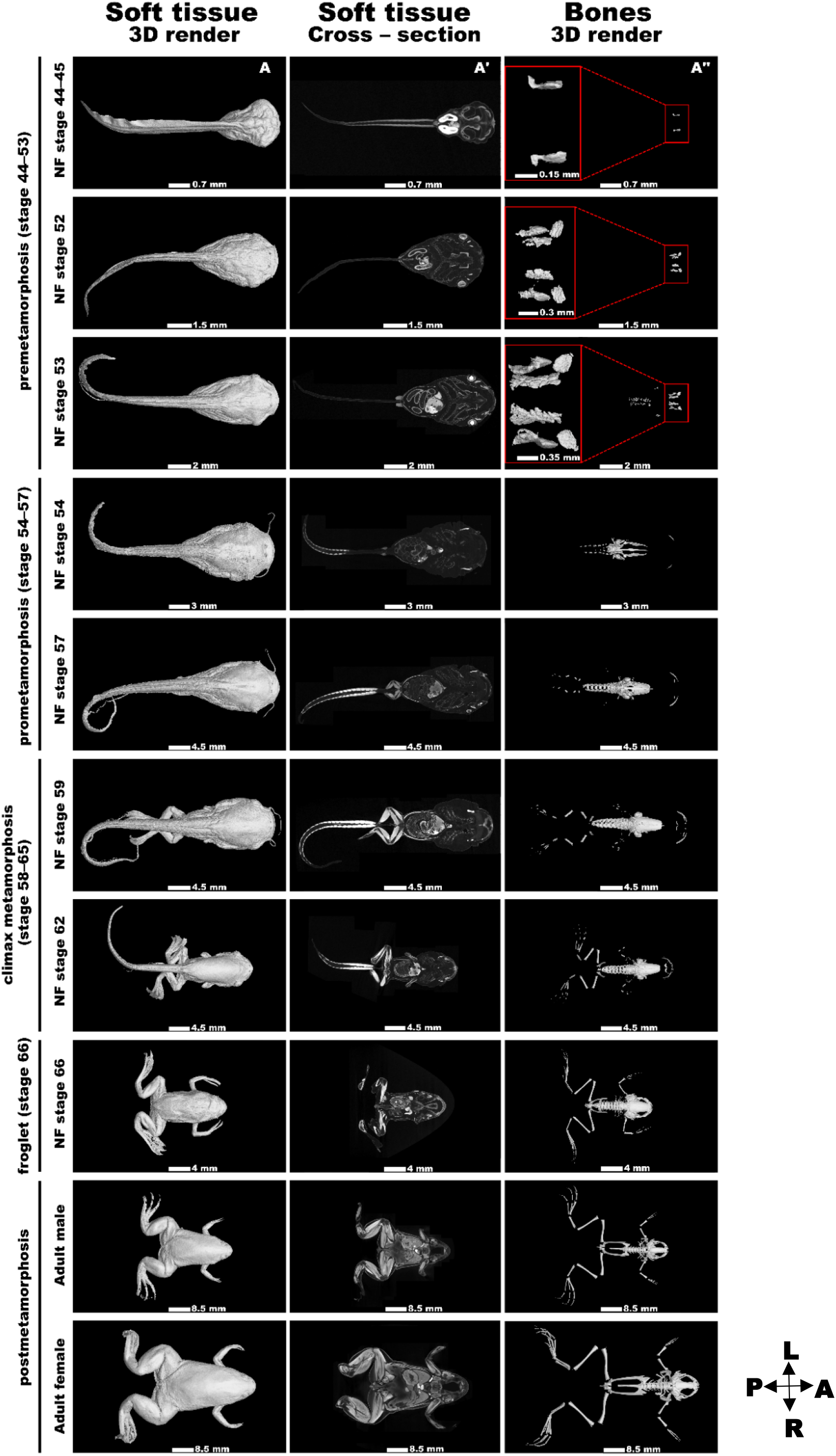
The atlas of *Xenopus laevis’s* late development. **A)** 3D renders of soft tissues of selected *Xenopus* developmental NF stages. **A’)** Cross-sections of soft tissues. The section level was selected based on capturing the brain area of *Xenopus* specimens. **A’’)** 3D renders of skeleton. In first three conditions (NF 44-53), the bones are zoomed in and shown in the red rectangle on left. The relevant scale bar is depicted on bottom in the center of each picture. All views are from dorsal view with cranial side pointing to the right. A: anterior; P: posterior; R: right; L: left.

### Head Development Analysis

First, we delved into the intriguing process of head development and skull metamorphosis in climax metamorphosis tadpoles, froglets, and adult frogs (refer to **Fig. 2A**; **Suppl. Video 1**). Our initial investigation employed morphometric analysis to shed light on the developmental changes in the skull. All skulls were oriented in the same direction (the anterior skull part heading to the top of a page; A: anterior; P: posterior; R: right; and L: left) in a top-bottom view. Such images can be further used, for example, to evaluate how the individual calvarian bones are being rearranged during metamorphosis or, for example, to shed light on facial region development.

**Figure 2.**
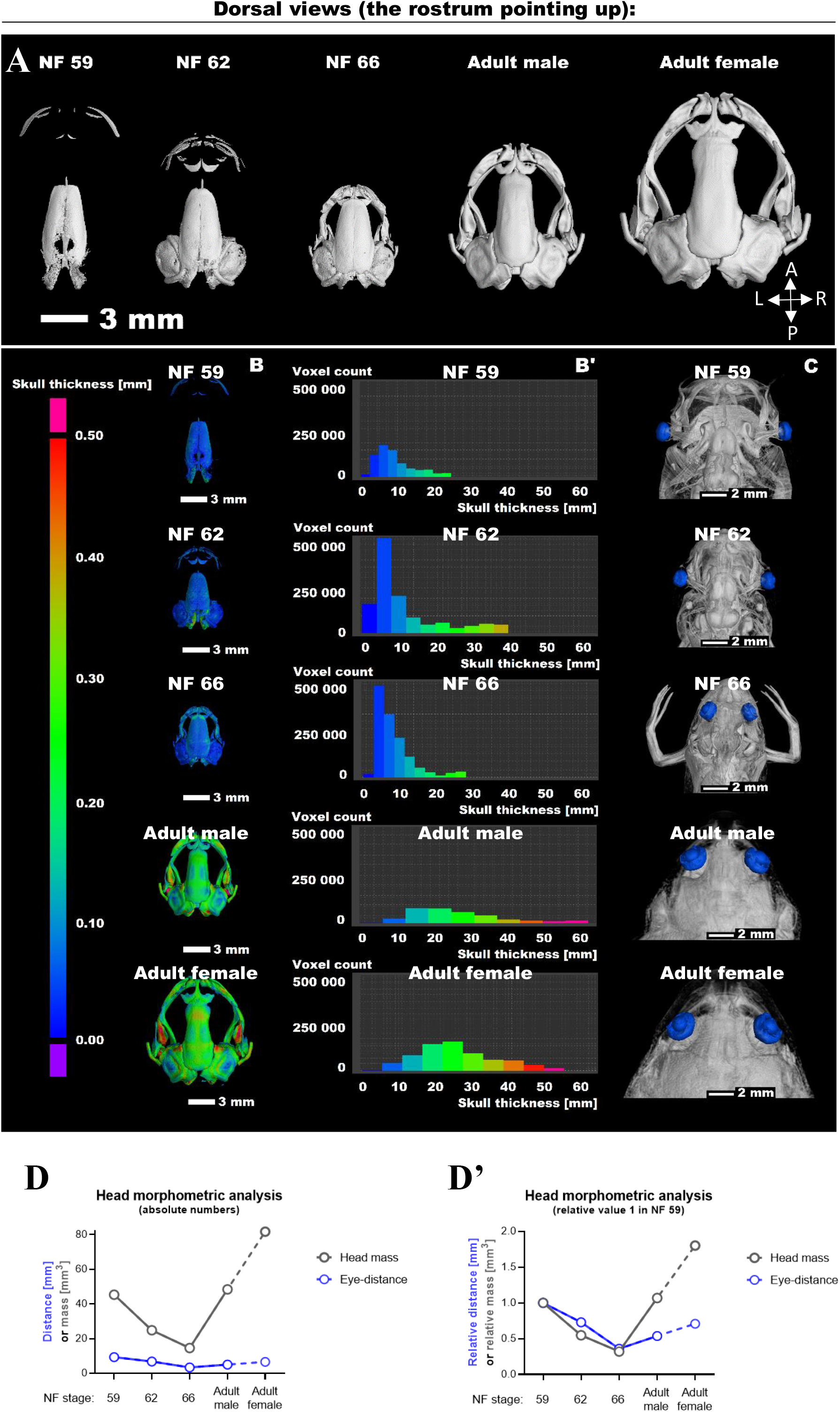
The analyses of the head development. **A)** The series of *Xenopus* skulls of selected stages. The scale bar with values in mm is shown on left bottom. **B)** The skull thickness of selected stages is shown. The scale bar with values in mm is shown on left side. B’) The relative distribution of bone thickness for each skull is shown. **C)** The overview of developing heads with highlighted eyes is shown. **D-D’)** Graphs displaying absolute and relative properties such as length in the head region such as eye distance (in mm), and head volumes (in mm^3^). In D’, the first value in NF stage 59 is normalized to 1.

Next, we performed the examination of skull thickness (**Fig. 2B-B’**; **Suppl. Video 2**). This analysis revealed a non-linear relationship, with adult frogs possessing significantly thicker skulls compared to their younger counterparts. This analysis suggests a positive correlation between skull mass and skull thickness.

Moving on, we made a noteworthy analysis regarding the eye distance during development, as illustrated in **Fig. 2C**. Our morphometric analysis (for an example of the measurement, see **Suppl. Fig. 2**) revealed a progressive decrease in eye distance during developmental stages (quantified in **Fig. 2D**). These changes actually occurred in *Xenopus* in two phases: a gradual increase in the pre-metamorphic stages (data not shown), followed by a peak at the onset of climax metamorphosis (NF stages 59 and 62), and finally, a decrease to a compact head configuration in froglets (NF stage 66) and adult frogs (**Fig. 2C**; quantified in **Fig. 2D**). This adaptation aligns well with the frog’s life strategy, transitioning from a water-dwelling tadpole with lateral eyes to an adult with eyes positioned on top of the head for a submerged lifestyle^30^, reminiscent of crocodilians^31^.

Based on the previous results, one can also explore the conservation analysis of the eye-head ratio with respect to frog gender. Our findings indicate that since one can quantify the mass volume precisely, female heads exhibit an increase in volume compared to males. However, the eye distance in females remains at only 78% of the head volume in males (see **Suppl. Table 3**). This observation may be linked to visual perception requirements, as the proximity of the eyes is still crucial for certain aspects of vision^32^. In summary, our micro-CT dataset can be further employed for the study of the head and its associated organs such as the eyes in *X. laevis*.

### Tooth Analysis

To enhance the depth of our analysis and increase its practical applicability, we conducted a comprehensive examination of the frog teeth, with an individual data set of an adult female jaw. Upon initial observation, the presence of teeth in *X. laevis* may not be readily apparent. This is due to the fact that teeth were exclusively found in the maxilla, referred to as maxillary teeth (**Fig. 3A-B**) and behind the maxillary arch on the vomeral bone – so-called vomeral teeth (not shown). Additionally, only a small portion of the maxillary tooth extended into the oral cavity^33^ (**Fig. 3C**). In contrast, the mandible contains no teeth (**Fig. 3D**). The shape of maxillary teeth is generally uniform (i.e., homodont dentition), typically resembling simple conical structures (see Fig. 3B). *Xenopus’* teeth exhibit an acrodont type of attachment, where they form an ankylotic attachment with the adjacent bone and are characteristic by regular renewal, a condition known as polyphyodont dentition. Notably, the phenomenon of tooth renewal in *Xenopus* is akin to what has been previously documented in other species, such as geckos^34,35^. In agreement with this, we present evidence of the replacement in the maxillary teeth (**Fig. 3E**). These analyses reveal the progression of ankylosis, from the early development of teeth located closer to the gingiva to their full fusion with the adjacent bone (see Fig. 3E). Our *Xenopus* micro-CT data set thus unveils the previously concealed dental structures of *X. laevis*, provides a high-resolution revelation of their “hidden” teeth, and offers new perspectives for studying teeth and their growth in *Xenopus*.

**Figure 3.**
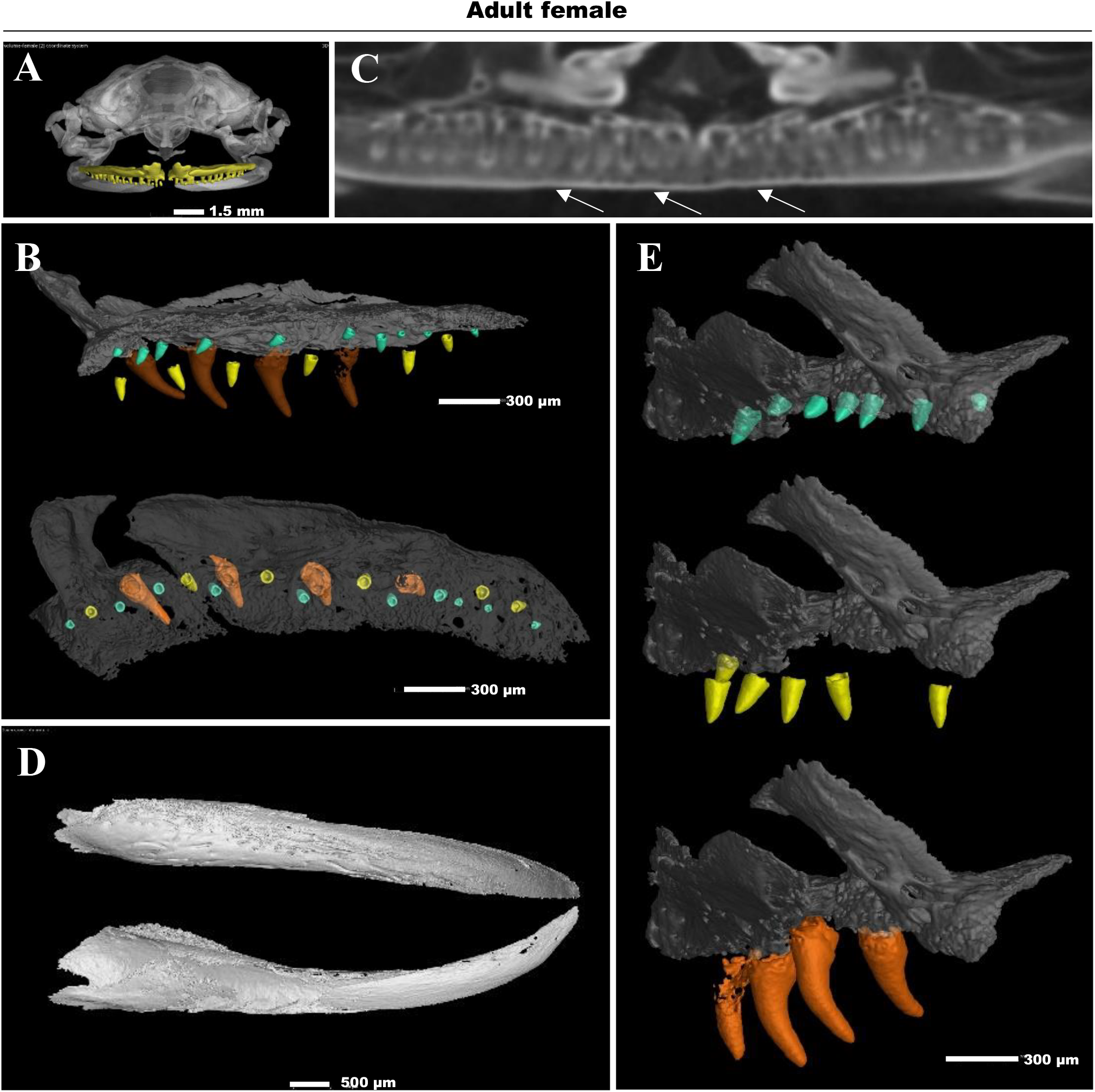
The analysis of teeth in an adult frog. **A)** The frontal view of an adult female frog skull is depicted with upper maxillary arch with maxillary teeth visualized in yellow. **B)** The lateral and top view on the right half of maxillary arch with visualized teeth from an adult female frog. Three different developmental stages of teeth are highlighted by different color (yellow, cyan, orange). **C)** The stained micro-CT scan shows that only a small portion of the tooth extends into the oral cavity. The arrows point on teeth rows which are not penetrated into oral cavity. **D)** The lateral view of mandible of an adult female frog confirms absence of teeth in this area. **E)** The lateral view of the rostral part of maxilla displays different stages of teeth during the replacement of tooth rows in detail.

### Micro-CT Data Analysis of Long Bone Growth Dynamics

Next, we assessed the growth dynamics of several long bones (shown in adult frogs in **Fig. 4A, B**; **Suppl. Video 3**; refer to **Suppl. Fig. 3** to see how the analysis was performed). Based on our micro-CT analysis, in terms of gender, we found that adult females can be viewed as essentially enlarged males, indicating a proportional growth pattern (**Fig. 4C**). We also investigated whether the development of the left and right sides of *X. laevis* is uniform or divergent (**Suppl. Fig. 4**), and consistently, we observed L-R length symmetry in all cases. Therefore, we next focused our detailed examination only on one side of the frog, specifically the left side (L).

**Figure 4.**
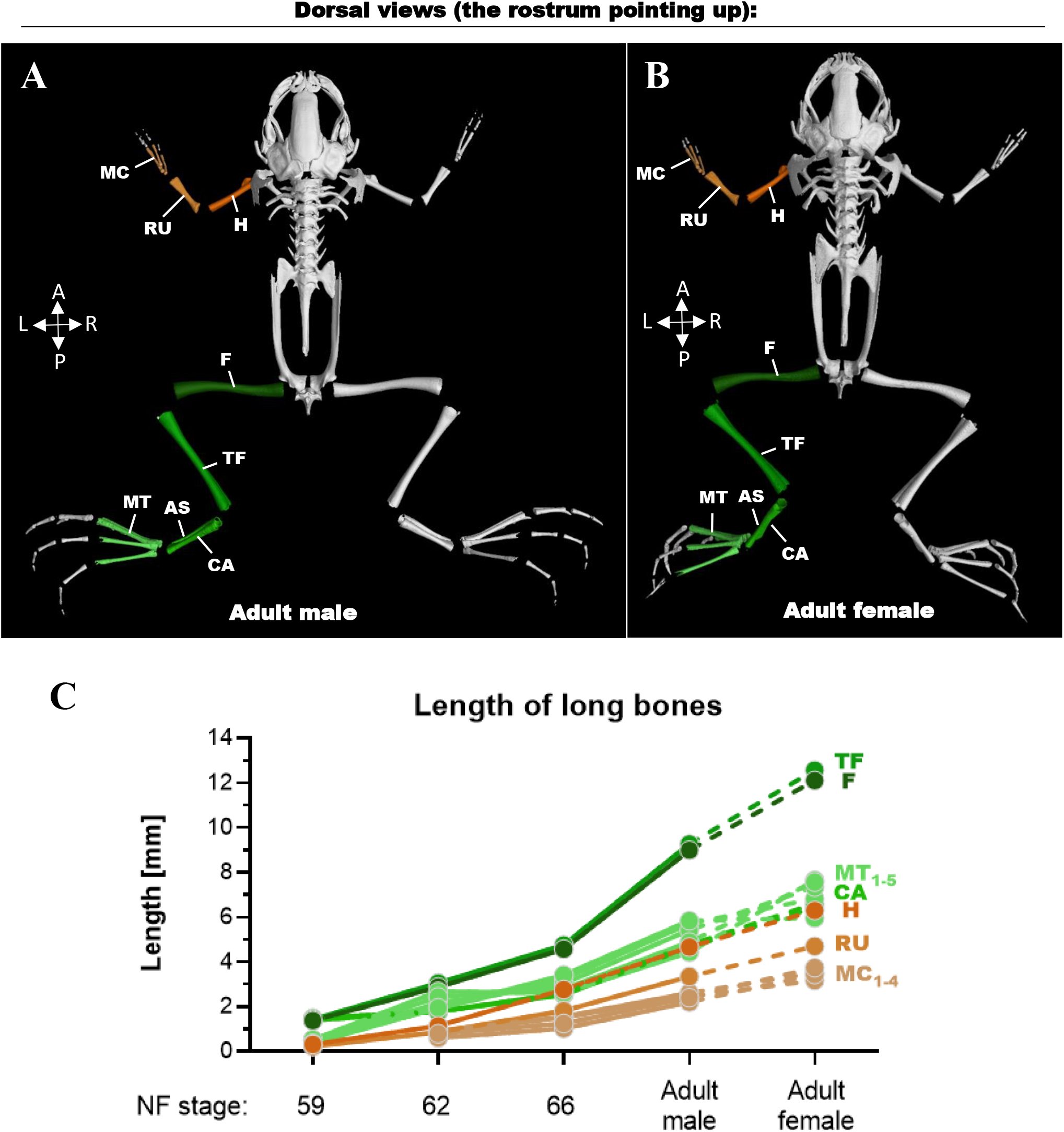
The analysis of frog skeletogenesis. **A)** The adult male frog with analyzed bones is depicted. **B)** The adult female frog with analyzed bones is displayed. The view of male and female is maximized and it is not proportional to each other. For their size proportions, see Fig. 1. **C)** The graph demonstrating the analyzed bones throughout the *Xenopus laevis’s* late development. A: anterior; AS: astragalus; CA: calcaneum; F: femur; H: humerus; L: left; MC: metacarpals; MT: metatarsals; P: posterior; R: right; RU: radioulna; TF: tibiofibular.

Our findings revealed that certain limb bones, particularly those in the hind legs, such as the femur and tibiofibular, exhibited relatively rapid growth, while other bones such as fore limb ones grew at a slower rate (**Fig. 4C**). Importantly, this difference was not attributed to absolute bone measurements but rather in relation to the animal’s overall length (data not shown). This phenomenon may be linked to the frogs’ utilization of their hind limbs for swimming, whereas even their forelimbs are primarily employed for food handling^36,37^. Moreover, the length of the tarsal bone such as astragalus and calcaneum appears to be less critical for escape or startle responses, such as swim kicking, compared to the lengths of the femur and tibiofibular, which are correlated with the muscle mass of the thigh and the calf. It is also intriguing to note that most bones initiate their growth from roughly the same size but end differently. For example, metatarsals develop at a faster rate than metacarpals (**Fig. 4C**), which aligns with functional considerations^37^. Together, based on our micro-CT data, one can support previously noted observations or make new predictions for further experimental testing of (long) bone growth.

### Ossification Analysis

Besides bone length, it is also feasible to analyze the bone mass using our *Xenopus* data set. Within the frog skeleton, two distinct types of bones can be generally distinguished in terms of their developmental origin. Dermal ossification, originating in the dermis, is evident in certain skull elements and two bones of the pectoral girdle, namely the cleithrum and clavicle. Conversely, the remaining components of the postcranial skeleton consist of either cartilage or cartilage-replacement bones, achieved through endochondral ossification^38^.

As the bones of the limbs undergo development in both tadpoles and adults, they typically comprise an ossified section alongside cartilage. Notably, the ossification process always commences at the central regions of long bones, as shown on the example of the femur (**Fig. 5A**). In assessing the mass ratio between cartilage and bone (**Fig. 5B-B’**), we observed that, with a few exceptions, all long bones share relatively similar proportions (data not shown). Furthermore, in both adult males and females, there is a lack of dynamic growth akin to the developmental stages, only a noticeable increase in mass (**Fig. 5C**). After climax metamorphosis, no proliferative ossification occurs, and the process is limited to the calcification of cartilage, irrespective of the frog’s gender (**Fig. 5C**). Moreover, the relative ratio of cartilage to ossified bone diminishes as development progresses, exemplified by the femur, a pattern consistent in other bones like the tibiofibular (data not shown). Thus, researchers can further take advantage of these indications to compare the growth of different bones in various aged animals.

**Figure 5.**
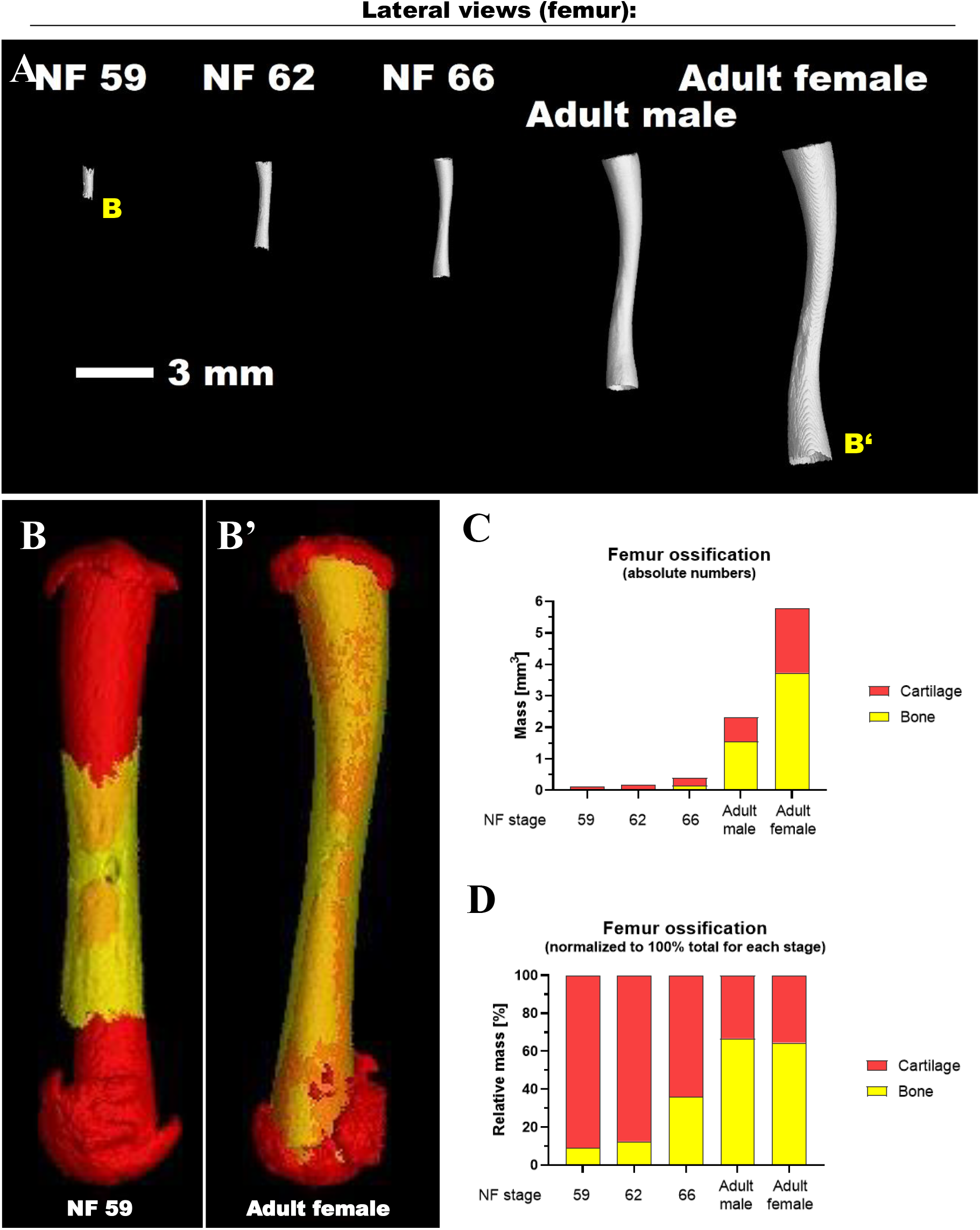
The analysis of longs bones including their cartilage and ossification. **A)** The collection of femur bone throughout the late development. **B-B’)** The femur from different developmental stages such as NF 59 (B) and an adult female (B’) with a focus on cartilage (in red) and bone (yellow). The view of bones is maximized and not in proportions. For size proportions, see Fig. 5A. The absolute (in **C**) and relative (in **D**) quantification of a femur bone and cartilage mass.P

### Segmentation of Selected Internal Soft Organ

Our dataset offers significant potential not only for the evaluation of hard bones but also for soft tissue such as internal organs. Precise segmentation of structures is essential because once the structure in focus is clearly distinguished, further assessment of its morphology and interspecies differences becomes considerably easier and more accurate. This tool also provides the flexibility to describe each internal structure in detail either separately or in the context of its individual surrounding elements.

In addition to studying differences during development, segmented structures can be utilized for investigating intersex variability. As an illustration of both phenomena, we focused on the key vertebrate organ, the brain, and observed the developing individual components of the tadpole and frog brain (see **Fig. 6A-E, A’-E’**; **Suppl. Video 4**). Subsequently, we also asked whether we could analyze nerves like *nervus opticus* using a micro-CT data set (**Fig. 6F**). We could see that the nerves attached to eyes slowly grew and then modified, as a specimen changed during development. In sum, researchers can now use our micro-CT dataset to explore internal organs such as the brain or structures such as nerves as well for further purposes.

**Figure 6.**
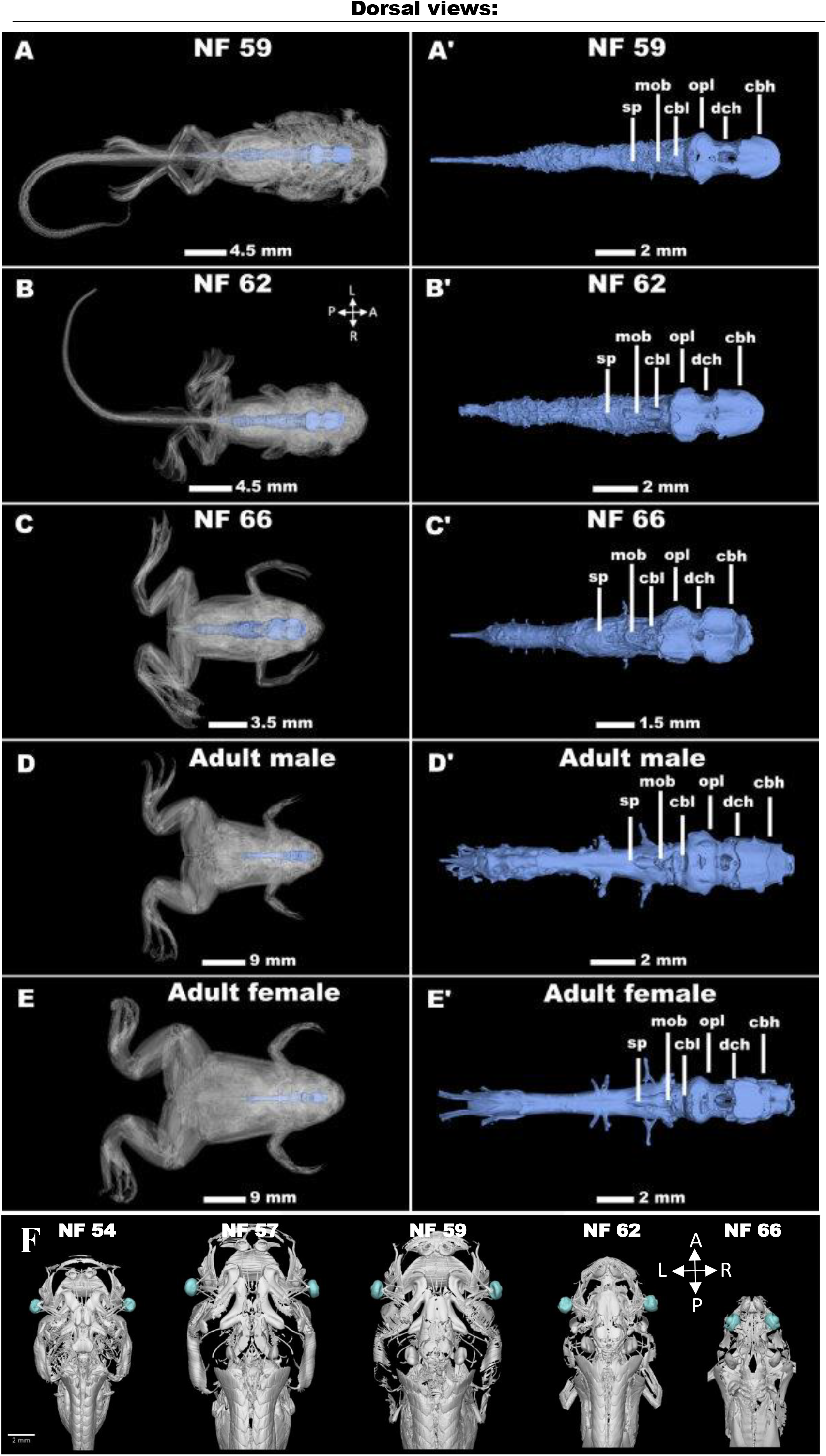
The analysis of brain development. **A-E)** Ind2iv2idual stages of *Xenopus* brain and their details in (A’-E’), the rostrum pointing to right, and **F)** nervus opticus attached to eyes (cyan), the rostrum is pointing to up. cbh: cerebral hemispheres; cbl: cerebellum; dch: diencephalon; mob: medulla oblongata; opl: optic lobes; sp: spinal cord.

### Further Biological Potential of *Xenopus’s* Data

The *X. laevis’* development dataset not only offers valuable insights into this amphibian species but also presents a wealth of biological potential as it serves as a starting point for comparative, regenerative, and evolutionary biology, and morphological anatomy. The dataset’s diverse species representation and the high quality of the scans open up a realm of possibilities for advanced subsequent analyses. One intriguing avenue of study lies in the examination of hard tissue growth. The dataset provides an ideal platform to investigate especially the long bones of *Xenopus* but also vertebrae, or a skull. Furthermore, the dataset facilitates the analysis of teeth or cartilage, which we have addressed using dedicated tools in VG Studio Max.

Furthermore, this dataset lends itself to in-depth exploration by researchers interested in the anatomy of *Xenopus* including internal organs. The dataset’s segmentation capabilities enable the precise isolation of individual structures, which can then be further processed to obtain detailed information regarding their shape, volume, and relationships with surrounding structures in three dimensions.

To enrich our understanding of adult *Xenopus* anatomy, morphology, and development, the dataset also includes scans of multiple *Xenopus* tadpoles. By combining these various analyses, the dataset offers comprehensive and detailed information on both hard and soft tissue morphology, making it a valuable resource for researchers studying this amphibian in the context of development.

In conclusion, this extensive dataset serves as a cornerstone for advancing our knowledge of amphibian morphological structures and evolutionary adaptations.

### Popularization/Educational Potential of Present Dataset

The *Xenopus* development dataset, which includes 3D representations such as views and videos of various developmental stages including the adult stage, holds significant educational potential. These 3D models can serve as invaluable learning tools for students at the university and secondary school levels worldwide, offering an innovative approach to understanding the complex process of *Xenopus* development. Additionally, these 3D models can be leveraged to engage the general public, for instance, in museums, providing an educational resource without the need for original specimens or samples (or their organs) preserved in fixatives.

The use of 3D scans for virtual reality applications further enhances the educational experience. It allows for a detailed exploration of individual structures and even provides the opportunity to virtually traverse through a *Xenopus* skeleton or explore specific external and internal organs. This immersive approach not only enhances the understanding of *Xenopus’s* development but also makes it accessible to a wider audience, fostering an appreciation for the intricacies of this fascinating organism.

## Data Availability

All volumetric data of the scanned *Xenopus* specimens can be accessed in the Zenodo repository (uploaded under link: 10.5281/zenodo.10214561). These datasets are presented as already-reconstructed 8-bit TIFF stacks. To enhance accessibility, the data was converted from its original 16-bit format to 8 bits, making it more compact for downloading and easier to open and analyze. This reduction in data size is particularly beneficial given the computational demands associated with working with large data sets.

The provided videos showcase individual animals and highlight analyses conducted using the scanned data, as detailed in the supplementary files. Additionally, .STL files accompany the image stacks, enabling users to visually inspect the scanned animals in three dimensions. These .STL models are also accessible via the Zenodo repository (10.5281/zenodo.10214561).

While the deposited .TIFF format data can be viewed in any basic image viewer, for a comprehensive exploration of the 3D nature of the images and the analyses performed, it is recommended to use a dedicated 3D data viewer. VG Studio MAX, a commercial software package from Volume Graphics GmbH, Germany, offers a wide range of tools for visualizing, manipulating, and analyzing 3D μCT image data. Its freeware version, MyVGL, is a suitable alternative for visualizing all datasets and their corresponding analyses. Another free software option is the Fiji ImageJ distribution from the National Institutes of Health, USA, capable of opening individual image stacks and creating 3D renders of all scanned *Xenopus* specimens.

In summary, the volumetric data of scanned animals, available on Zenodo repository, provides efficient accessibility through 8-bit TIFF stacks, complemented by .STL models, with recommended 3D exploration using software like VG Studio MAX, MyVGL, or Fiji ImageJ.

## Additional Files

**Supplementary Video 1:** The skull of an adult *Xenopus* female in 3D view.

**Supplementary Video 2:** The skull of an adult *Xenopus* female in 3D view while using wall thickness analyses.

**Supplementary Video 3:** The skeleton of an adult *Xenopus* female in 3D view.

**Supplementary Video 4:** The brain of an adult *Xenopus* female in 3D view.

## Abbreviations

A: anterior
AS: astragalus
CA: calcaneum
cbh: cerebral hemispheres
cbl: cerebellum
dch: diencephalon
F: femur
H: humerus
L: left
MC: metacarpals
micro-CT: micro-computed tomography
MMR: Marc’s Modified Ringers
MT: metatarsals
mob: medulla oblongata
NF stage: Nieuwkoop and Faber stage
opl: optic lobes
P: posterior
R: right
RU: radioulna
sp: spinal cord
SS: suprascapular
TF: tibiofibular.

## Authors’ contributions

J.H. conceived the research idea, manipulated with *Xenopus*, wrote the draft, and supervised the overall work.

J.L. analyzed the data and contributed to the draft writing, data visualization, and figure creation.

M.K. conducted micro-CT experiments and analyzed the data.

A.R., P.R.H., J.Kr., M.B., and J.H. provided expertise in the biological interpretation of the data. J.Kr., T.Z., J.Ka., M.B., and J.H. secured funding for the project.

M.K., M.B., and J.H. developed the initial concept of the *Xenopus* atlas. All authors edited and approved the manuscript.

