## Supplementary materials for "Unveiling Vertebrate Development Dynamics in Frog *Xenopus laevis* using Micro-CT Imaging"

These Supplementary materials contain 4 Figures and 3 Tables.

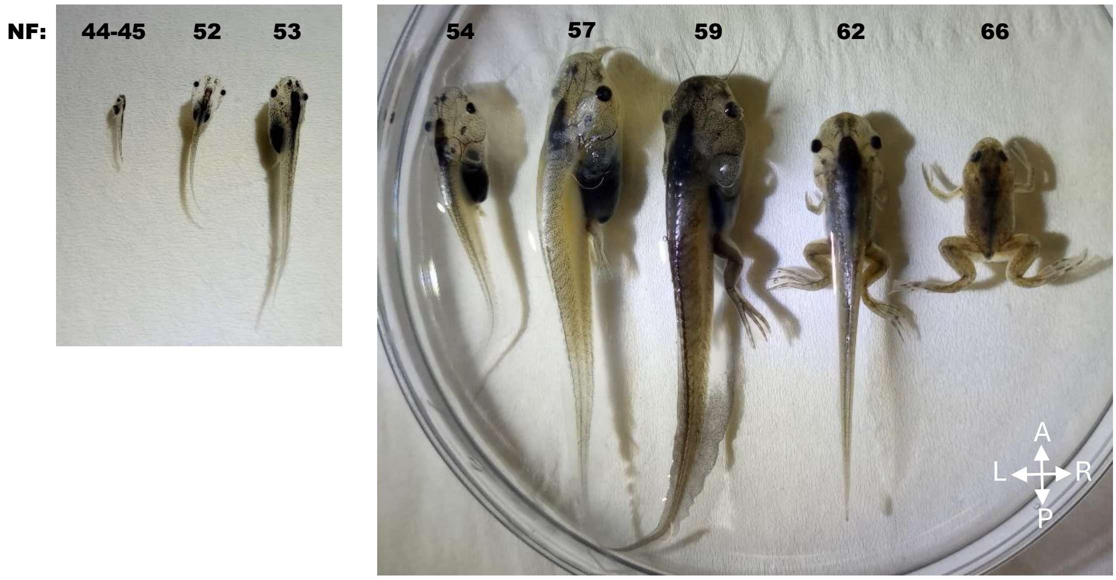


**Supplementary Figure 1.** The macroscopic view of tadpoles used for micro-CT analysis. Numbers above the organism indicate the corresponding NF stages.

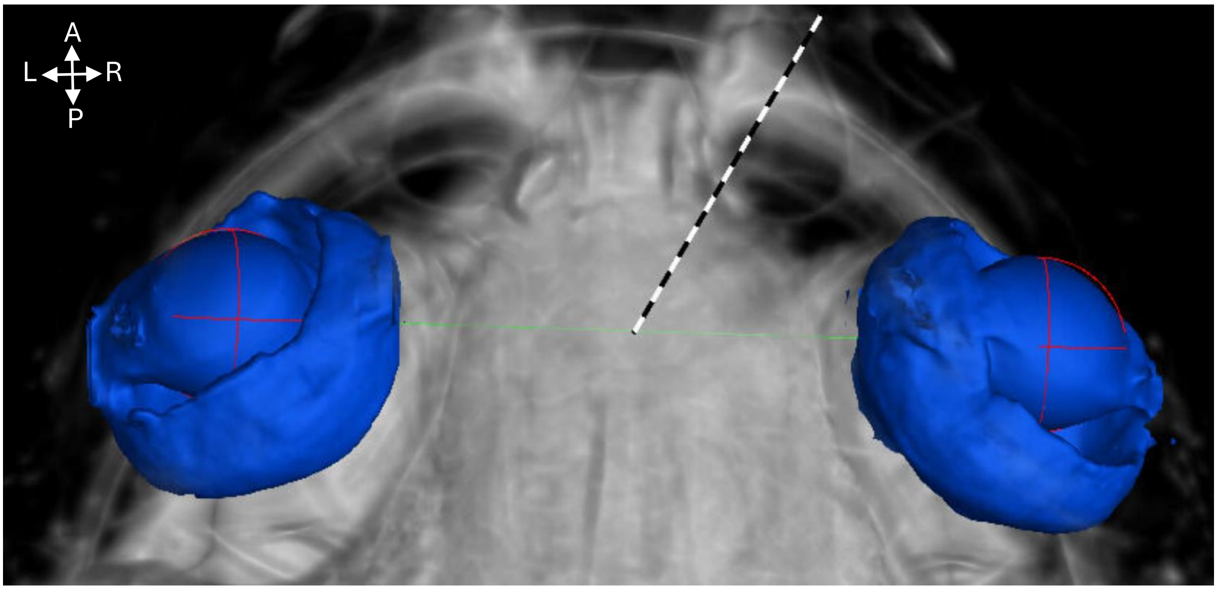


**Supplementary Figure 2.** The representation of how the eye distance was calculated. The green line is the distance between two eyes marked with red balls on left and right. The example with adult male frog is shown.

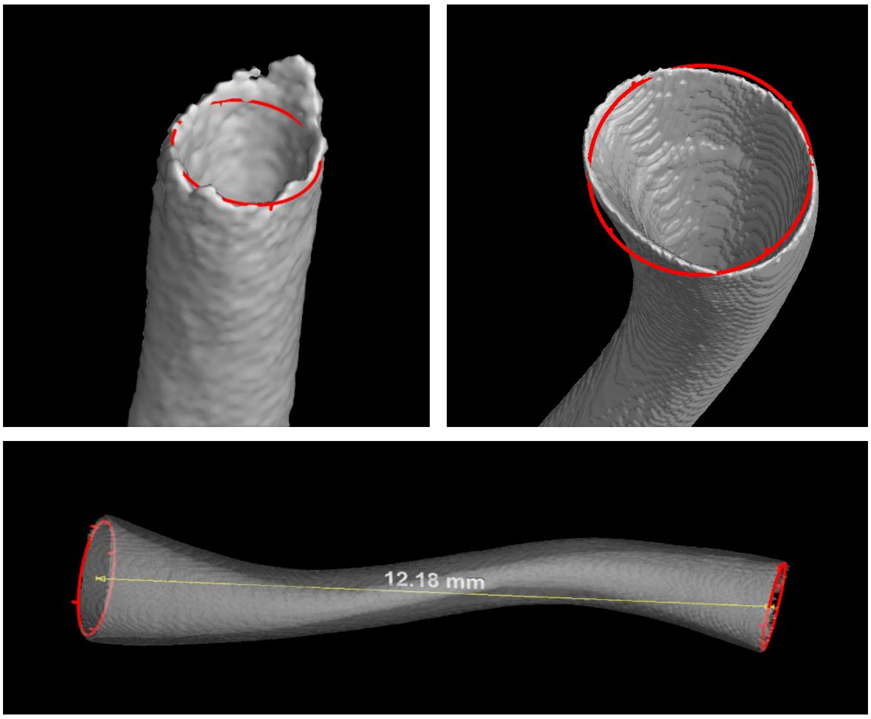


**Supplementary Figure 3.** The diameter analysis method used for long bone measurement. The red circle inserted to the end of a bone served as a reference for the analysis of bone lengths. The circle was used for the labelling of the one side of the bone (top). The distance between two circles was measured to determine the bone length (below).

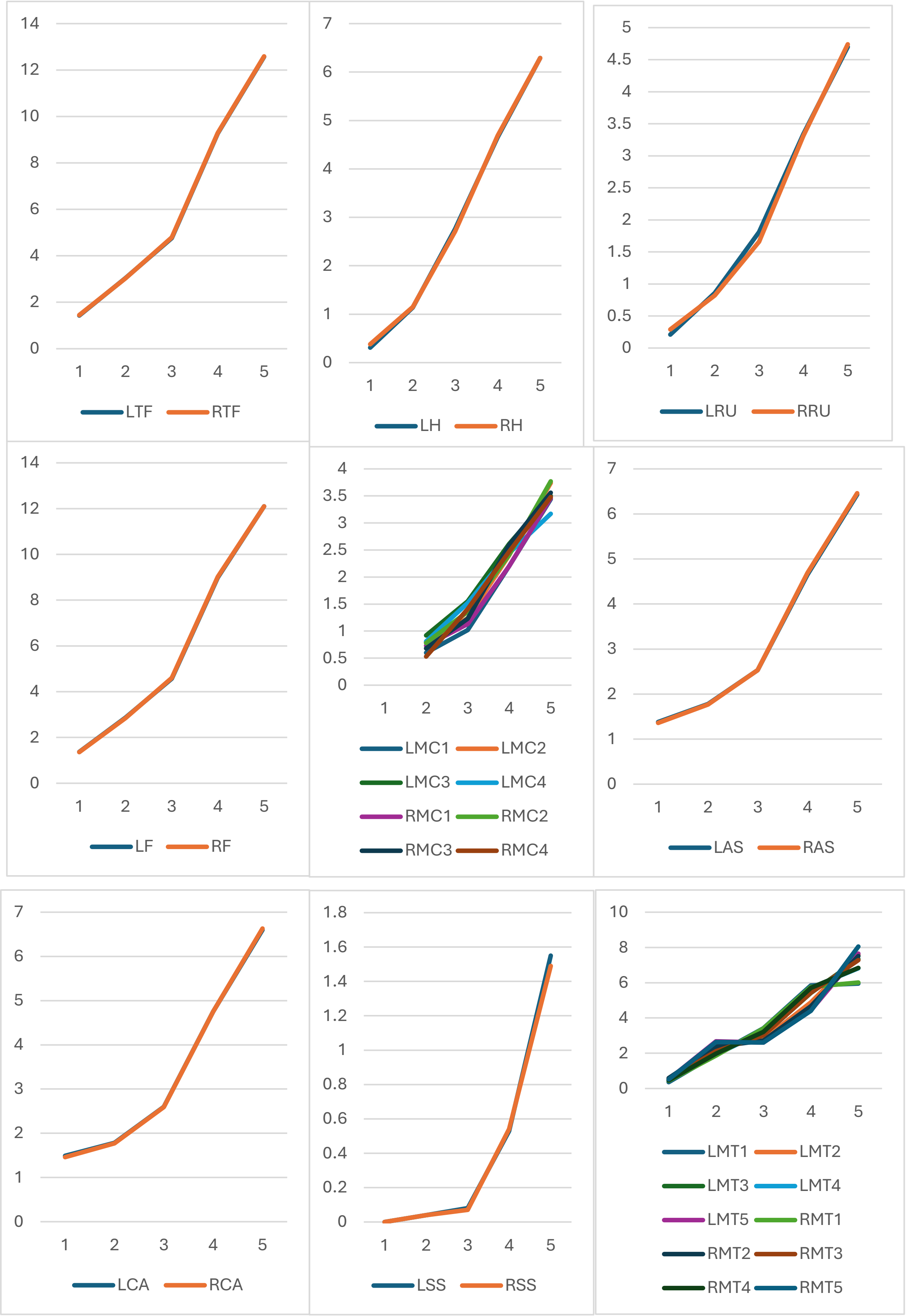
LTF RTF
LH RH
LRU RRU
LF RF
LMC1 LMC2 LMC3 LMC4 RMC1 RMC2 RMC3 RMC4
LAS RAS
LCA RCA
LSS RSS
LMT1 LMT2 LMT3 LMT4 LMT5 RMT1 RMT2 RMT3 RMT4 RMT5

**Supplementary Table 1. The detailed procedure of micro-CT scanning.**

| sample | voltage [kV] | current [μA] | timing | binning | images | filter (mm) Al | scan time | resolution [μm] | multiscan |
|---|---|---|---|---|---|---|---|---|---|
| Premetamorphosis_tadpoles | 60 | 200 | 400 | 2 | 2000 | 0.2 | 56 min | 5 | 3 tiles |
| Premetamorphosis_tadpoles_ stained | 60 | 200 | 600 | 2 | 2000 | 0.2 | 3h 30 min | 5 | 3 tiles |
| Metamorphosis_stages | 60 | 200 | 500 | 2 | 2600 | 0.3 | 4h 30min | 15 | 3 tiles |
| Metamorphosis_stages_stained | 60 | 200 | 500 | 2 | 2600 | 0.3 | 4h 30min | 15 | 3 tiles |
| Frog_adults | 60 | 200 | 500 | 2 | 2200 | 0.3 | 1h 30min | 40 | |
| Frog_adults stained | 80 | 200 | 500 | 2 | 2200 | 0.2 | 1h 15min | 40 | |

**Supplementary Table 2. The detailed protocol of soft tissue staining.**

| | NF stage 44-45 | NF stage 52 | NF stage 53 | NF stage 54 | NF stage 57 | NF stage 59 | NF stage 62 | NF stage 66 | adults |
|---|---|---|---|---|---|---|---|---|---|
| EtOH 30% | 1 h | 4 h | 8 h | 8 h | 12 h | 12 h | 12 h | 8 h | 24 h |
| EtOH 50% | 1 h | 4 h | 8 h | 8 h | 12 h | 12 h | 12 h | 8 h | 24 h |
| EtOH 70% | 1 h | 4 h | 8 h | 8 h | 12 h | 12 h | 12 h | 8 h | 24 h |
| EtOH 80% | 1 h | 4 h | 8 h | 8 h | 12 h | 12 h | 12 h | 8 h | 24 h |
| EtOH 90% | 1 h | 4 h | 8 h | 8 h | 12 h | 12 h | 12 h | 8 h | 24 h |
| 1% $I_2$ MeOH 90% | 1 h | 4 h | 12 h | 20 h | 26h | 26h | 26 h | 22h | 2d + 2d + 4d |
| EtOH 80% | 20 min | 30 min | 1 h | 1 h | 1 h | 1 h | 1 h | 1 h | 2 h |
| EtOH 50% | 20 min | 30 min | 1 h | 1 h | 1 h | 1 h | 1 h | 1 h | 2 h |
| EtOH 20% | 20 min | 30 min | 1 h | 1 h | 1 h | 1 h | 1 h | 1 h | 2 h |

**Supplementary Table 3. The eye distance to head mass ratio in *Xenopus* displays gender-related differences.**

| | Adult male | Adult female |
|---|---|---|
| Eye distance [mm] | 5.00 | 6.61 |
| Head mass [$mm^3$] | 48.43 | 81.71 |
| The ratio of eye distance to head mass | 0.103 | 0.081 |
| **Normalized to 100% for male** | **100 %** | **78 %** |
